# Facilitating Gene Editing in Human Lymphoma Cells Using Murine Ecotropic γ-Retroviruses

**DOI:** 10.1101/2024.12.03.626526

**Authors:** Manish Kumar, Eva Gentner-Göbel, Palash Chandra Maity

## Abstract

Genetic modifications using CRISPR-Cas9 have revolutionized cancer research and other pre-clinical studies. Exceptionally, these efficient tools are inadequate in a few disease models and cell lines due to the aberrant differentiation states and the accumulation of excessive somatic mutations that compromise the robustness of viral gene delivery and stable transduction. A couple of B lymphoma cell lines fall into this category where lentiviral transfection becomes inefficient and exhibits variable efficiency. Additionally, lentiviral delivery requires high biosafety levels. To address this challenge, we have developed a two-step strategy that supports CRISPR-Cas9 through lentivirus and murine ecotropic γ-retrovirus. By engineering B lymphoma cell lines to express Cas9 and mCat-1, a specific receptor for ecotropic retroviruses, we enable efficient and safe gene editing through ecotropic γ-retrovirus. We demonstrate the efficacy of this method by generating IgM-deficient B lymphoma cell lines. This innovative approach simplifies protocols, enhances accessibility, and paves the way for standardized gene manipulation of B cell lymphoma models for molecular cell biology research.

## 1. Introduction

The cell and molecular biology research, including preclinical studies on therapeutic biologicals, rely on three major factors: the availability of robustly maintainable cell lines, high-efficiency gene delivery methods, and precision tools like CRISPR-Cas9 for genome editing. Alongside these technical considerations, there are variable levels of biosafety issues associated with the entire process, specifically concerning viral transduction-mediated gene delivery and CRISPR-Cas9 protocols [1–3]. There is a continuous effort to establish a standard molecular biology and translational research protocols that allow gene manipulations without necessitating higher biosafety levels and other Genetically Modified Organism (GMO) requirements [4,2]. In this context, we explore the possibility of using CRISPR-Cas9 genome editing and exogenous overexpression in B lymphoma cell lines rendered permissive to murine ecotropic γ-retroviruses.

For mature B cell lymphoproliferative diseases and normal B cell signaling studies, there is a spectrum of B lymphoma cell lines available, categorized according to disease of origin and relevant subtypes such as Diffuse Large B Cell Lymphoma (DLBCL) or Waldenström macroglobulinemia (WM), also known as Lymphoplasmacytic Lymphoma (LPL). The WM cells exhibit mixed characteristics resembling both plasma cells-like antibody secretion and lymphoid-like membrane-bound antibody expression as B cell receptor (BCR). Despite having a large catalog of B cell lines originating from leukemia and lymphoma conditions, the use of specific cell type with the correct BCR isotypes and other coreceptor expressions impose limitations on many *in vitro* study designs. In most cases, BCR isotype expression and other immune receptors’ expression become skewed and altered in stable cell lines compared to *in vivo* conditions. One way to circumvent this situation is through exogenous overexpression of receptors via high-efficiency gene delivery by viral transduction.

Due to the terminal state of differentiation and accumulation of somatic mutations, most primary malignant B cells as well as lymphoma-derived cell lines are resistant to stable transductions [5,6]. Transductions with standard amphitropic retroviruses and pantropic viruses like Vesicular stomatitis virus G protein (VSV-G) pseudotyped lentivirus often yield unsatisfactory transduction efficiencies [7,5]. Recently, lentivirus systems pseudotyped with either Gibbon Ape Leukemia Virus (GaLV) envelope or Feline Leukemia Virus envelopes (FeLV) were developed to enhance transduction efficiencies for B cell research [8,6]. Both amphitropic retroviruses and VSV-G pseudotyped lentiviral protocols require at least biosafety laboratory 2 (BSL-2) environment and are subjected to GMO regulations [9,10]. Modifications like GaLV or FeLV pseudotyped lentiviruses for improved viral transduction efficiency necessitate further stringent biosafety laboratory environments due to relatively high safety concerns. These regulations may differ among different institutional and regional biosafety authorities. A safer alternative to these pantropic lentivirus systems and amphitropic retroviruses are ecotropic retroviruses. Normally, ecotropic retroviruses can only infect rodent cells as they enter cells through specific receptors, namely murine cationic amino acid transporter-1 (mCat-1) or murine solute carrier family 7 member 1 (Slc7a1) receptor [1,11]. The mCat-1 protein is ubiquitously expressed in all the murine cell lines, especially in B cells and all hematopoietic-origin cell types. Bringing mCat-1 expression into the human cells renders them permissive to ecotropic retroviral transduction.

CRISPR-based genome editing technology is an indispensable tool for almost every genetic modification, ranging from double-stranded break and subsequent gene silencing to programmable epigenome engineering in combinations with transcriptional activators or repressors [12]. For genome editing, the CRISPR system functions as a bipartite system consisting of a single-stranded guide RNA (sgRNA) and the Cas9 endonuclease. The system can be implemented either as a two-vector system or a single plasmid driving sgRNA expression under U6 promoter and Cas9 expression under CMV-like promoter. Modifications and tagging of Cas9 enzymes with other functional domains alter the specificity and outcome of the specific CRISPR method. Combining CRISPR-based genome editing and viral gene delivery methods, we manipulate BCR expression for molecular oncology research in B lymphoma cell lines. Due to the large size of Cas9 cDNA and obligatory additional modifications that together approaches the packaging limit, the two-vector system is often preferable for viral transductions. In contrast, a single plasmid carrying both sgRNA and Cas9 cassettes is used for liposome and electroporation-based transfection protocols. In best practice for gene knockout, we generate a Cas9-expressing stable cell line by lentiviral transduction followed by the introduction of specific sgRNA using different alternative methods of transient or stable transfections. For molecular oncology studies, transient transfection of sgRNA is preferred as the construct is lost and it avoids permanent sgRNA expression. Therefore, restoration of gene function by subsequent expression of cDNA requires no sequence adjustment at the sgRNA target site.

While using a two-vector system or a single plasmid successfully produces a knockout cell line, a second transduction for restoring the gene expression is relatively difficult in B lymphoma cell lines [13]. Similarly, researchers wish to avoid the production of pantropic viruses encoding both Cas9 and sgRNA due to higher biosafety levels and other safety issues. To overcome these issues, we first generate a lymphoma cell line stably expressing Cas9 protein. Additionally, these Cas9-positive cells co-express mCat-1 protein necessary for ecotropic γ-retrovirus transduction, which can be performed at standard BSL-1 environment. We designed a lentiviral construct for co-expressing Cas9 and mCat-1. Thus, upon the selection of stably expressing cells free from lentivirus-vector particles, these cells can be maintained and engineered by ecotropic γ-retrovirus transduction at BSL-1 conditions.

Here, we provide a robust protocol for using the CRISPR–Cas9 genome editing system in combination with ecotropic retrovirus transductions (Fig. 1). Using this method, we demonstrate the deletion of BCR isotype IgM in two different lymphoma cell lines, namely, SU-DHL-6 and BCWM.1. The protocol demonstrates a step-by-step guide to lentiviral transduction for the generation of Cas9-expressing ecotropic retrovirus-permissive (Cas9^+^Eco) lymphoma cell lines, followed by ecotropic retrovirus particle-mediated sgRNA expression and screening of IgM-deleted clones.

**Figure 1.**
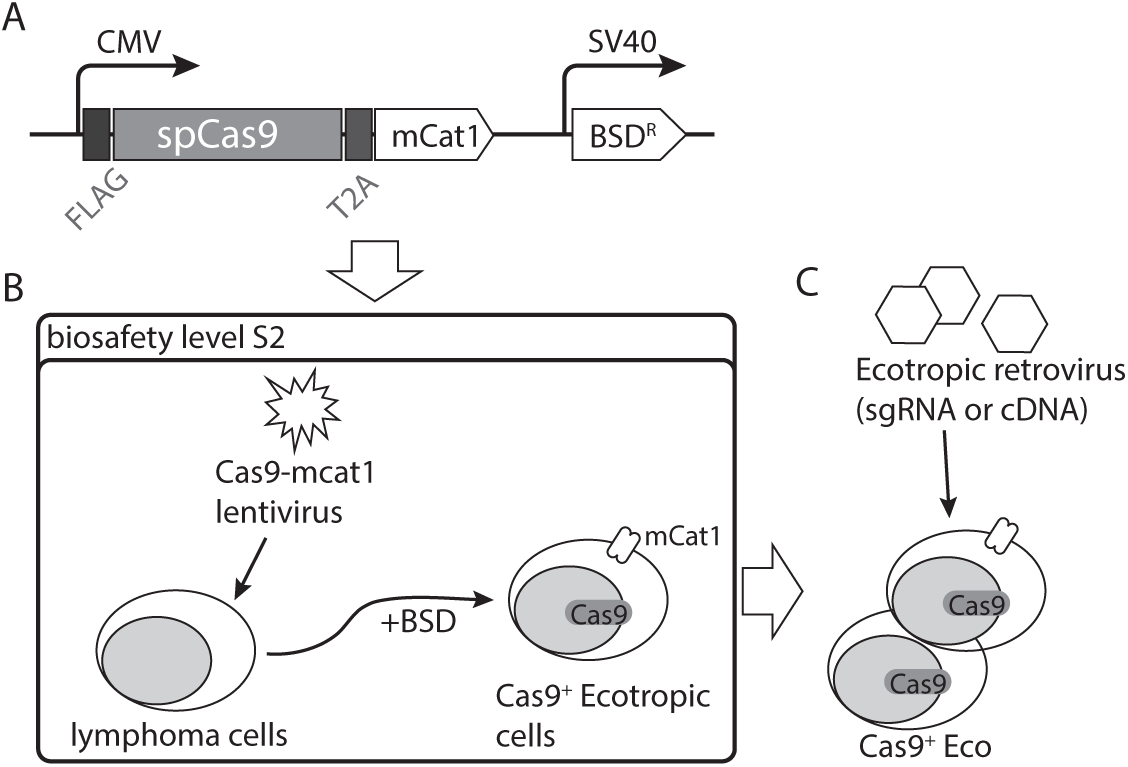
Schematic overview of the method. A. Design and generation of lentiviral plasmid expressing N-terminal FLAG-tagged spCas9 followed by self-cleaving T2A motif fused to receptor for γ-Retroviruses, murine cationic amino acid transporter-1 (mCat-1), all under the control of MSCV promoter. Additionally, the plasmid expresses Blasticidin resistance (BSD^R^) under the control of SV40 promoter placed at the 3’ of Cas9-T2A-mCat-1 cassette. The lentiviral vector backbone pCDH was used to generate the plasmid pCDH MSCV Cas9-T2A-mCat-1 BSD^R^ (in short pLCas9-mCat1 BSD) B. Preparation of lentiviral particle in Lenti-X™ 293T packaging cell line and infection of human B lymphoma cell lines followed by two weeks of Blasticidin selection to generate Cas9 and mCat-1 expressing ecotropic γ-Retrovirus permissive Cas9^+^ eco-tropic cell lines. The entire process of lentivirus generation till completion of Blasticidin selection must be carried out at biosafety level 2. Upon removal of viral supernatant and multiple washing steps and media exchanges for another two weeks during Blasticidin selection, cells which are free from any residual viral particles can be confirmed by performing PCR reaction from the culture media. C. The Cas9^+^Eco cell lines can now be maintained and transduced with murine ecotropic γ-Retroviruses under normal cell culture conditions.

## 2. Materials

### 2.1. Cell lines

1. SU-DHL-6: B-cell lymphoma (DSMZ Repository ID: ACC-572) originated from a non-Hodgkin Lymphoma patient with peritoneal effusion assigned to germinal center B-cell-like (GCB) DLBCL. This cell line expresses IgM(IgD)/Igκ B-cell receptor (BCR).
2. BCWM.1: B-cell LPL or WM (Cellosaurus Repository ID BCWM.1 CVCL_A035) cell line derived from bone marrow aspirate derived from Waldenström’s macroglobulinemia patient [14]. BCWM.1 is a widely used model for IgM secreting LPL/WM. This cell line expresses BCR subtype IgM(IgD)/Ιgλ.
3. Lenti-X™ 293T: This cell line is a sub-clone of human embryonic kidney (HEK) cells for lentivirus particle production (Takara Bioscience). It delivers high transfection efficiency and supports the expression of viral proteins.
4. HEK 293T or its derivative Phoenix-Eco (ATCC Repository ID: CRL-3214) cells is required for packaging of the retrovirus with ecotropic helper plasmids.

### 2.2. Plasmids and oligonucleotides

1. pL Cas9-mCat-1 BSB plasmid generation: This lentiviral construct is designed to express N-terminal FLAG-tagged spCas9 and murine cationic amino acid transporter-1, mCat-1 (Fig. 1). The plasmid was constructed by using pCDH_MSCV (System Bioscience) vector backbone carrying spCas9 cDNA from Addgene #48138 and mCat-1 cDNA from Addgene #17224 (see Note 1).
2. Lentivirus packaging helper plasmids: pMD2.G, Addgene #12259 and pxPAX2, Addgene #12260 were used for VSV-G pseudo typed lentiviral packaging.
3. Retroviral sgRNA expression plasmid generation: pRsgRNA IGHM (cµ1) mRFP1 was derived from Retro-gRNA-mRFP1, Addgene #112914. It encodes for sgRNA targeting 5’ of human IGHM gene constant domain exon 1 Cµ1 (see Note 2). Targeting sequence with probable cut site (^) is 5’ CCC GTC GGA TAC GAG CA^G CG 3’. Primer pair for amplifying ∼500bp region encompassing the sgRNA target site is Fwd: 5’ CAA AGA CCT GAG GCC TCA CCA CGG CC 3’ and Rev: 5’ GCT GGA CTT TGC ACA CCA CGT GTT CG 3’.
4. Retroviral packaging helper plasmid: pCL-Eco, Addgene #12371 is used for efficient packaging of ecotropic retroviruses.
5. Retroviral expression vector: pMIG (derived from Addgene #9044) is used as retroviral expression vector.

### 2.3. Cell culture

1. Complete DMEM for Lenti-X™ 293T and HEK293T cell lines: Dulbecco’s Modified Eagle Medium (DMEM) containing high glucose, supplemented with 2 mM L-Alanyl-L-glutamine solution (stable glutamine), 100 U/mL Penicillin and 100 µg/mL Streptomycin, 10 mM HEPES and 10% heat-inactivated (hi) fetal bovine serum (FBS) of south American origin. Phoenix-Eco cells are cultured in Iscoves’s modified DMEM (IMDM) completed with stable glutamine, Pen-Strep, HEPES and hiFBS. Cells cultured in complete IMDM is better maintained in 7.5% CO2 incubator instead of standard 5 % CO2 condition.
2. Complete RPMI for B cell lines: Roswell Park Memorial Institute (RPMI) 1640 media supplemented with 2 mM stable glutamine, 100 U/mL Penicillin and 100 µg/mL Streptomycin, 10 mM HEPES, 1 mM sodium pyruvate, 50 µm β-mercaptoethanol and 10 % hiFBS.
3. Cell culture grade phosphate buffer saline (PBS), pH 7-7.2.
4. General cell culture material: Adherent type 10 cm Petri dish and 6-well plate, suspension type 12-well-plate, T-25 and T-75 tissue culture flasks, 0.45 µm syringe-mounted filters and 3 mL Luer Lock syringe.

### 2.4. Transfection and transduction

1. Lentivirus packaging and transduction: Transfection medium PolyFect (Qiagen), Lenti-X™ Concentrator (Takara Bio), 10 mg/mL Polybrene, 1 mg/mL RetroNectin® (Takara Bio) and 20 mg/mL Blasticidin. All procedure related to lentiviral packaging and transductions until complete removal of viral particles were carried out in BSL-2 environment and approved by institutional genetic engineering and GMO regulatory authority (See Note 3).
2. Retroviral packaging and transduction: Transfection medium GeneJuice® (Millipore) and 10 mg/mL Polybrene.
3. PES membrane-based Ultrafiltration device with 10-20 kDa cut-off fitted inside a 15 mL tube. Amicon® Ultra-4 centrifugal unit (Millipore) or Vivaspin® centrifugal concentrator (Sartorius).

### 2.5. Antibodies and immunochemicals

1. anti-mouse mCat-1 (clone no. SA191A10) APC conjugated.
2. anti-FLAG (clone no. M2) Alexa Fluor 647 conjugated.
3. anti-IgM (clone no. MHM-88) PerCP-Cy5.5 conjugated.
4. FIX&PERM® cell fixation and permeabilization kit (Nordic MUbio) for intracellular staining.

### 2.6. Equipments

1. Biosafety level 2 (BSL-2) laboratory environment equipped with CO2 incubator and benchtop centrifuge
2. PCR instrument and molecular biology working set up.
3. Fluorescence activated cell (FACS) analyzer and sorter with appropriate laser and filter set up.

## 3. Methods

### 3.1. Generation of Cas9 expressing lymphoma cell line

#### 3.1.1. Preparation of lentiviral particles

The entire procedure of lentiviral packaging and infection to the lymphoma cell lines until complete removal of viral particles is carried out in BSL-2 environment and approved by genetic engineering and GMO regulatory authority (See **Note 3**).

1. Virus-producing Lenti-XTM 293T cells were seeded in complete DMEM one day before their transfection in order to reach a confluency of 70% on the day of transfection. Keep the number of cells within the range of 300-500K cells per well (∼9.6cm2) of a 6-well plate giving a spreading density of 30-50,000 cells per cm2. Keep the total volume of complete DMEM medium within 2-3 mL per well of the 6-well plate. For scale up of the entire protocol to generate higher volume of virus, larger sized plates can be optimized (see <u>Note 4</u>).
2. For preparation of the transfection mixture for 1 well (9.6 cm2) of 6-well plate, take 1.8 µg pLenti-cas9-mCat-1-BSD target plasmid, 1.4 µg psPAX2 and 0.8 µg pMD2.G helper plasmids all in a 1.5 mL tube, add up to 150 µl serum-antibiotic free DMEM media, and mix well by vortexing. Bring down the liquid at the bottom of the tube by brief centrifugation.
3. Add transfection reagent, 15 µl PolyFect and mix immediately by vortexing briefly for 5-10 sec (see <u>Note 4</u>). Try to bring down the liquid at the bottom of the tube by finger tapping (avoid centrifugation) and incubate for 15 min at room temperature (RT).
4. To determine the optimum amount of plasmid required for each transfection, we varied the total amount of plasmid keeping a fixed ratio of the pLenti-cas9-mCat-1-BSD, psPAX2 and pMD2.G at 9:7:4 according to their molecular weight (Fig. 2A). To estimate the percent of transfected Lenti-XTM cells, intracellular FACS staining is performed after taking out the virus supernatant (see <u>Note 5</u>).
5. Meanwhile, gently remove all the media, wash with 1 ml of DMEM and add 1 ml of fresh complete DMEM media. While washing and changing media from multiple wells at a time, care should be taken to avoid drying of seeded wells.
6. Add 350 µL of complete DMEM to the transfection mixture tube, mix by pipetting 2-3 times up and down and overlay this 500 µL on virus-producing cells (Lenti-XTM 293T or 293T cells) drop-wise with 150 µl of transfection mixture. Gently swirl the plate to ensure uniform distribution of the complexes.
7. Cells are then incubated at 37°C in a humidified 5% CO2 incubator for 48-72 hours. After 72 hours viral supernatant or virus containing medium (VCM) is collected and filtered through a 0.45 µm syringe-mounted filter. VCM can be used directly, stored at 4°C for 1-3 days or kept at -80°C for long-term storage. It is recommended to store VCM at -80°C (see <u>Note 2</u>.). As per the regular biosafety guidelines, the entire process of lentivirus VCM production, usage, and storage is restricted to S2 laboratories (see <u>Note 6</u>).

**Figure 2.**
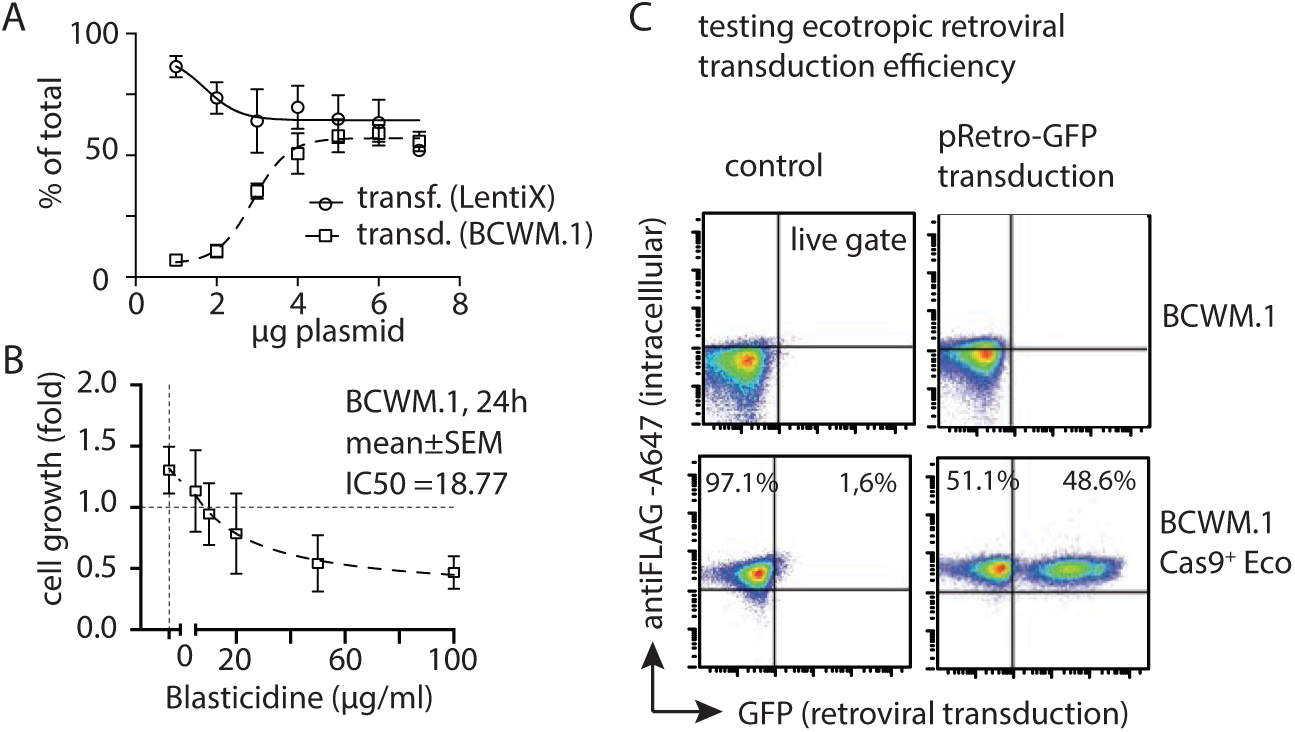
Generation and testing of ecotropic γ-retrovirus permissive B lymphoma cell line. A. Optimization of lentiviral particle preparation and transduction efficiency. Different amounts of GFP reporter lentiviral plasmid pCDH MSCV_EF1-copGFP-Puro were used together with helper plasmids pMD2.G and psPAX2 at a ratio of 1.5:1:1.5. Transfection efficiencies of the Lenti-X™ 293T cell line at different amounts of plasmids were measured by GFP positive cells in FACS analyses upon PFA fixation after 48 hours. The corresponding transduction efficiency of lymphoma cell line BCWM.1 using the generated viral supernatant were determined and plotted. B. Optimization of Blasticidin dose on BCWM.1 cell line. Fold change in cell growth was determined by analyzing the live cell count after 24 hours of Blasticidin treatment at different doses. C. Representative FACS plots show Cas9 expression and ecotropic retrovirus infection measured by intracellular staining of FLAG-tag and GFP expression, respectively. The BCWM.1 untreated (upper panels) and BCWM.1 Cas9Eco (lower panels) cells were transduced with retroviral particles generated from pRetro-IRES-GFP empty vector and analyzed by FACS after 5 days of infection.

#### 3.1.2. Optional preparation of lentiviral concentrate and long-term freezing

1. After collecting the filtered VCM through 0.45 µm pore syringe-mounted filter, collect the whole 9 mL of supernatant from wells of the 6-well plate, add 3 mL Lenti-X™ Concentrator and mix well by inverting the tubes. Incubate mixture at 4°C for 30 minutes to overnight (O/N) in refrigerator (see Note 6).
2. After incubation, centrifuge sample at a speed of 1,500 x g for 45 min at 4°C. After centrifugation, an off-white pellet will be visible. Carefully remove supernatant, taking care not to disturb the pellet. Leave approximately 0.5 mL liquid at the bottom to ensure no disturbance to the thick slurry containing the concentrated virus pellet.
3. Add 3.5 mL of serum-free RPMI to the 0.5 mL pellet and mix well. Add this 4 mL sample to the PES membrane-based ultrafiltration device with 10-20 kDa cut-off fitted inside a 15 mL tube. Spin the device at 3,000 x g for approximately 10–20 min in a 4°C precooled swing-bucket rotor. After complete filtration it leaves approximately 200-500 μL depending on the density of particles.
4. To wash the concentrator chemicals from the viral particles, add up to 4 mL of serum-free RPMI media into the filtration unit. Remove the filtered liquid from the bottom prior to starting the centrifuge. Repeat one more time washing by adding up to 4 mL of serum-free RPMI media on the concentrator. After the centrifugation, take out 200-400 µL of remaining VCM concentrate from the bottom of the filtration unit using a 200 μL tip. Collect VCM concentrate the in a 2.0 mL freezing vial and add complete RPMI media up to 1.25 mL. VCM concentrate can be stored in -80°C for future use (see Note 6).

#### 3.1.3. Lentiviral transduction and generation of the Cas9-mCat-1 expressing cells

1. One day before performing the transductions, each well of 6-well plate was coated with 1 mL of 20-50 µg/mL RetroNectin® (stock 1 mg/mL) in PBS. Preferably, use adherent type of 6-well plate. Incubate plates at room temperature (RT) for 30 minutes or at 4°C for 16-18 hours till ON in a refrigerator. If using 50 µg/mL RetroNectin®, it can be collected and stored at -20°C for the second use. It is important to keep a control well left untreated for RetroNectin® coating, instead containing equal volume of PBS. This control well will receive target cells but no virus, therefore can be left untreated for sustainability and saving of expensive RetroNectin® reagent (see <u>Note 7</u>).
2. On the day of transduction, remove the RetroNectin® coating solution and add 1 mL of 2% BSA in each well for 30 min at RT or until adding virus supernatant and cells.
3. Add 2.5 mL VCM (∼2 wells of 6-well plate) or up to 1.25 mL concentrated VCM (∼all 6 wells of 6-well plate) depending on the desired multiplicity of infection (MOI) (see Note 8). Prior to adding on to the RetroNectin®-coated plate, all VCM is supplemented with 8 µg/mL Polybrene (from 10 mg/mL stock). Thereafter, plates are incubated and centrifuged at 1800 rpm for 3 hours at 32-37°C. It is important to keep a no-virus control well containing equal volume of complete RPMI instead of VCM.
4. Discard 2.25 mL and 1.0 mL of the supernatant from VCM and concentrated VCM-coated plates, respectively. Optional, unbound virus particles can be removed by washing each well with 1 mL of serum-free RPMI media. To avoid drying of the plates, discard the washed supernatant leaving 250-500 µL media just before adding the cells.
5. While centrifuging the plates for virus coating, prepare the target cells in appropriate volume of complete RPMI giving 500K cells/mL. Remove the washing supernatant from the virus-bound plate and immediately add 2 mL of target cells (BCWM.1 or DHL-6) in each well of the 6-well plate at a density of 1×105 cells/cm2. Add equal number of cells in no-virus control well, required to measuring transduction efficiencies (see Note 5).
6. To maximize the contact between the target cells and viral particles, plates can be centrifuged at 1200 rpm for 30 min at 32-37°C. Without disturbing the settled cells, transfer the plates in a humidified 37℃, 5% CO2 incubator for 2 days.

#### 3.1.4. Selection of Cas9^+^Eco cell lines

1. After two days, observe the cell growth pattern and compare growth rate by visual inspection in comparison to no-virus control. Collect all the adherent and non-adherent cells in 15 mL tubes, centrifuge to remove supernatant and wash the cell pellet with 5 mL of complete RPMI medium. Depending on the growth rate of the transduced cells compared to no-virus control, target cells can be diluted in larger volume at an estimated seeding density of 100-250K cells/mL and cultured in larger vessels or flasks.
2. Repeat step 1) every two days until 6-8 days depending on the cell growth. Now there are several millions of cells to start the selection against the BlasticidinTM resistance (BSDR). Before starting the selection procedure, transduction efficiency is measured by intracellular FACS staining with anti-FLAG antibody. Transduction efficiency is plotted as function of variable plasmid input and equivalent amount of VCM, (see Note 5). Since the required dose of antibiotic varies largely between cell types, it is necessary to determine the BSD sensitivity of the target cell lines and culture condition. Our optimized BSD dose is 20 mg/mL (Fig. 2B). It is preferable to perform the selection procedure in 6-well plates for easy medium change without centrifugation and washing of cells until the resistant clones start proliferating and repopulating.
3. Plate the cells in 6-well plates at a dilution of 1×106 transduced cells in 3 mL of complete RPMI medium added with 20 µg/mL of BSD. In parallel to transduced cells treated with no-virus control cells with the same dose of BSD.
4. After two days observe the cells under the microscope and compare the transduced clones versus no-virus control for the presence of bright living and dead cell fractions. Remove 1-2 mL of the culture from the top without disturbing the settled cells at the bottom and add equal volume of complete RPMI containing 20 µg/mL of BSD. If necessary, a small aliquot of 100 µL cells can be taken out for determining the live-dead fractions by FACS analyses upon 1 µM Sytox Blue staining (See Note 5).
5. Repeat step 4) for 2-4 times until the resistant clones start proliferating and repopulating. Once the resistant clones start growing, wash the cells by centrifugation and plate them in larger vessels at a density of 100-250K cells/mL.
6. At this stage, cells can be taken to normal BSL-1 environment after analysing the culture supernatant for the presence of lentiviral particles either by performing capsid protein ELISA or qPCR analyses [15]. Usually after two weeks of culturing the transduced cells no traces of active viral particles are remaining.
7. Positively selected cells can be tested for mCat-1 expression and Cas9 expression by performing either western blot analysis or FACS staining. To detect the 2xFLAG-tagged Cas9 expression, we perform intracellular staining with anti-FLAG Alexa Fluor 647 using FIX&PERM® cell fixation and permeabilization kit according to manufacturer’s instruction. Similarly, perform FACS analyses for mCat-1 expression using anti-mouse mCat-1 APC antibody. For some unknown reason, often mCat-1 expression is undetected by surface staining, but nicely detected by intracellular FACS staining.

### 3.2. Generation of IgM knock-out by retroviral transduction of Cas9^+^Eco lymphoma cell lines

#### 3.2.1. Preparation of retroviral particles and transduction

The entire procedure of retroviral packaging and infection of the Cas9^+^Eco lymphoma cell lines are carried out in BSL-1 environment and approved by institutional genetic engineering and GMO regulatory authority (See **Note 3**).

1. Retrovirus producing Phoenix-Eco cells were seeded in complete IMDM (see <u>Note 9</u>) one day before their transfection to reach a confluency of 70% on the day of transfection. Keep the number of cells within the range of 400-500K cells per well (∼9.6cm2) for a 6-well plate giving a spreading density of 40-50,000 cells per cm2. Keep the total volume of complete DMEM medium within 2-3 mL per well of the 6-well plate. For scale up of the entire protocol to generate higher volume of virus use large sized plates.
2. For preparation of the transfection mixture for 1 well (9.6 cm2) of 6-well plates mix 1.5 µg pRsgRNA IGHM (cµ1) mRFP1 target plasmid and 0.5 µg pCL-Eco helper plasmid all in a 1.5 mL tube, add up to 100 µl serum-antibiotic and other additive-free IMDM media, and mix well by vortexing. For testing the retroviral transfection efficiency, use 1.2 µg of pMIG plasmid expressing GFP. Bring down the liquid at the bottom of the tube by brief centrifugation. Add transfection reagent, 4 µl of GeneJuice® and mix immediately by vortexing briefly for 5-10 sec. Try to bring down the liquid at the bottom of the tube by finger tapping (avoid centrifugation) and incubate for 15 min at room temperature (RT).
3. Meanwhile, gently remove all the media, wash with 1 mL of complete IMDM and add 1 mL of fresh complete IMDM media. While washing and changing media from multiple wells at a time, care should be taken to avoid drying of seeded wells.
4. Add 400 µL of complete IMDM to the transfection mixture tube, mix by pipetting one time up and down and overlay this 500 µL drop-wise onto Phoenix-Eco cells. Gently swirl the plate to ensure uniform distribution of the complexes.
5. Cells are then incubated at 37°C in a humidified 7.5% CO2 incubator for 48-72 hours. After 72 hours viral supernatant or retrovirus containing medium (rVCM) is collected and filtered through a 0.45 µm syringe-mounted filter. The rVCM can be used directly or stored at 4°C for 1-3 days. Unfortunately, concentrating retroviruses and long-term storage of these ecotropic retroviruses at -80°C do not work efficiently, at least in our hands, and are therefore not recommended. Optional, retroviruses can be concentrated, and media exchanged from IMDM to RPMI by using Retro-X™ Concentrator (see Note 6).
6. One day before transduction, Cas9+Eco cells can be splitted and kept at 250-500K cells/mL seeding density to obtain the robust exponential growth phase on the day of transduction. If retrovirus concentration and media exchange is wished, collect the whole 9 mL rVCM from a 6-well plate in a 15 mL tube and add 3 mL of Retro-X™ Concentrator, mix well and incubate O/N at 4°C. Next day, centrifuge sample at 1,500 x g for 45 min at 4°C and resuspend the pellet in 1 mL of complete RPMI.
7. On the day of transduction, harvest Cas9+Eco cells and prepare a stock of 2×106 cells/mL in complete medium. For transduction, use 250 µL (500K) cells and 150 µL (300K) cells in each well of 12- and 24-well plate, respectively. Optionally, plates can be pre-coated with 0.1 mg/mL of sterile poly-D-Lysine solution for 30 min at RT and then remove excess of poly-D-Lysine by washing the wells 2 times with distilled water and one time final wash with PBS (see Notes 7). Poly-D-Lysine coating enhances the settling of the B cells and reduces the time of spin inoculation. Coated plates must be used on the same day within few hours of coating.
8. Add 8 µg/mL of polybrene to rVCM and apply 1 mL or 0.5 mL of polybrene supplemented rVCM to each well of 12- or 24-well plates, respectively. It is important to keep a control well receiving only cells and equivalent amount of complete RPMI but no rVCM. While using concentrated stock, equivalent amount of rVCM can be diluted in the complete RPMI and supplemented with 8 µg/mL of polybrene prior to applying to the cells. Plates are then centrifuged at 1,200 rpm for 90 min at 32-37°C in swing-bucket rotor plate centrifuge. After centrifugation, plates are incubated at 37°C in 5% CO2 incubator for another 2 hours. When using poly-D-Lysine coated plates, 5 min centrifugation is sufficient to bring down the cells. Thereafter, plates are incubated for 4 hours in the cell culture incubator prior to media change.
9. After incubation with rVCM, take off and discard the supernatant carefully from the top without disturbing the settled cells. Avoid complete removal of the media to keep the cells undisturbed. Approximately, 1 mL and 0.5 mL media removal per well is sufficient for 12- and 24-well plates, respectively. In this case the poly-D-Lysine coating helps retaining the settled cells. Add 1 mL and 0.5 mL of fresh complete RPMI media per well for 12- and 24-well plates, respectively. Plates are then incubated in 5% CO2 cell culture incubator for O/N.
10. Next day, repeat step 9), exchange 1 mL and 0.5 mL media per well of 12- and 24-well plates, respectively. Plates are then incubated in 5% CO2 cell culture incubator for 2 more days.
11. After 72 hours of transduction, collect all the adherent and non-adherent cells in 15 mL tubes, centrifuge to remove supernatant and wash the cell pellet with 5 mL of complete RPMI medium. Depending on the growth rate of the transduced cells compared to novirus control (from step 8), target cells can be diluted in larger volume at an estimated seeding density of 100-250K cells/mL and cultured in larger vessels or flasks.
12. Repeat step 11) every two days for 2-3x times, giving final wash on 6-8 days. Now there are several millions of cells to start the sorting for single cell cloning or selection against the antibiotic resistance, if any.

#### 3.2.2. Testing retroviral transduction and generation of single cell IgM deleted clones

1. For testing the efficacy of mCat-1 and retroviral transduction, we transduce Cas9+Eco cells with empty vector retroviral construct pMIG expressing GFP reporter (Fig. 2C). Gene expression can be determined after 4-5 days of retroviral transduction. In this case, both Cas9+Eco BCWM.1 and WT BCWM.1 cells are transduced with either pMIG rVCM or no-virus control. After 5 days post transductions percent of GFP positive cells are analyzed by FACS upon intracellular staining with anti-FLAG Alexa Fluor 647 (Fig. 2C).
2. For the efficacy of sgRNA targeting IgM first exon, cells were analyzed by FACS for mRFP1 expression and surface IgM BCR level at day 5, 7 and 10 post transduction (Fig. 3A). Loss of IgM expression is visible in both Cas9 Eco DHL-6 and Cas9 Eco BCWM.1 cells.
3. To obtain the single cell clones, IgM negative and mRFP1 positive cells are sorted into a U-bottom 96-well cell culture plate at a rate of 1-3 cells per well (see Note 10). To avoid drying of the sorted droplet, 50 µL of complete RPMI media were added to each well of the plates prior to sorting. Plates are incubated in 5% CO2 cell culture incubator for 3-5 days and replenish with another 100 µL of complete media.
4. After 1 week of sorting, plates are monitored every 2-3 days to assess possible contamination and appearance of the clonal expansion of cells in individual wells. Number of growing wells can be recorded as a function of days post sorting and number of cells per well. To avoid minimal contact and loss of clonogenic potential of a single cells, it is advisable to sort 1 cell per well for 5-10 plates. In addition, 2-3 plates each with 2 and 3 cells per well. Upon growing, cells can be further sorted as single cell to ensure monoclonality.
5. Most of single cell colonies, if survive, grow within 3-4 weeks. However, it is not expected to find all the growing clones at the same time because of their differential growth rate. Collect cells from the growing wells time to time and plate them in larger wells stepwise from 96-well to 48-, 24-, 12- and 6-well plates.
6. After 4 weeks, as sufficient number of growing colonies are collected, analyze the individual clones by FACS staining for mRFP1 expression and surface IgM BCR level as mentioned before. Thereafter, IgM negative clones were selected for further analyses to determine the clonality and the pattern of mutation introduced.
7. Isolate genomic DNA from the IgM negative clones, and amplify the approximately 400-500 bp target region encompassing the sgRNA site. In this case, a 484 bp fragment is PCR amplified using the specific primer pair described in section 2.2 Plasmids and oligonucleotides.
8. Fragments are either gel eluted or directly purified from the PCR rection after confirming the amplification of the 500 bp fragment in sample volume. Afterwards, purified PCR products are analyzed by Sanger sequencing using the Fwd amplification primer. A clear unambiguous sequence spectra on both side of the sgRNA target site represents single cell colony (see Note 10). Upon aligning to the IGHM genomic DNA encompassing the target region, determine the actual mutation or deletion resulting aberrant stop codon causing the loss-of-function mutation in IgM (Fig. 3B).

## 4. Notes

1. The Cas9-mCat-1 lentiviral construct is generated in two step process. At first, pCDH_MSCV vector is linearized by EcoRI and DraIII and used as a vector backbone to perform Gibson assembly with amplified spCas9 cDNA and mCat-1 cDNAs connected through a self-cleaving p2a peptide sequence. Second, the resulting plasmid is linearized by XmaI and SalI and assembled with BSD expression cassette under the control of SV40 promoter. In parallel to wild type spCas9 that causes double strand breaks, the alternative nickase version of mutant spCas9 D10 is available in the same lentiviral format. The nickase spCas9 facilitates highly specific gene editing by reducing the off-target effect using a pair of sgRNA targeting adjacent region shorter than 20-30 bp.
2. The sgRNA sequences targeting IGHM Cµ1 exon are obtained from CRISPR site finding tool available to Geneious Prime® 2023 software using the previously described methods [16,17]. It encodes for sgRNA targeting 5’ of human IGHM gene constant domain exon 1 Cµ1.
3. It is mandatory to perform all procedure related to lentiviral production in BSL-2 environment. These include stages from transfection of plasmids in the packaging cell lines until completion of transduction and complete removal of viral particles (approximately two weeks) confirmed by ELISA or qPCR analysis [15]. In addition, an approval from institutional genetic engineering and GMO regulatory commission or relevant authority is required.
4. There are various alternative methods and transfection reagents including lipid or synthetic peptide-based compounds available for putting viral DNA into the packaging cell lines. We demonstrate the method with PolyFect (Qiagen) reagent. It can be adapted to other transfection reagents by optimizing the transfection and transduction efficiency. For example, using in-house prepared stock of transfection grade linear Polyethylenimine Hydrochloride (MW 40,000), we find similar results at 2.5-fold higher amount of total plasmid per well (9.6 cm2) of a 6-well plates. Notably, the increase in total amount of plasmid as input in trafection sometimes causes decreation in trasfection efficiency measured by the reporter gene experssion, in this case intarcellular anti-FLAG staining (Fig. 2A). However the transduction efficieny of the produced viral supernantat will increase. This is because of the combination of target and helper plasmids and optimum ratio or copy numbers of target and helper plasmids in maximum number of packaging cell lines. While optimizing the amount of plasmid, it is important to maintain the ratio of target and helper plasmids constant, in this case 9:7:4, to ensure proportional copy numbers based on their individual molecular weight. Similarly, we only demonstrate the lentiviral preparation using Lenti-XTM 293T packaging system, which can be replaced by other HEK293T derived cell lines used for mammalian protein expression.
5. Instead of optimizing the transfection and transduction efficiency using a general reporter construct expressing GFP or similar, we directly use the Cas9-mCat-1 plasmid. This is to avoid the discrepancy between the reporter-based optimization values and the transfection of actual large size lentiviral plasmid. Therefore, we suggest using the actual target plasmid to optimize the protocol. To determine the transfection efficiency and transduction efficiency, we use intracellular FACS staining to detect the FLAG-tagged Cas9 expression or mCat-1 expression. We demonstrate the protocol by using FIX&PERM® kit, which can be replaced by in-house protocol and reagents such as 4% Paraformaldehyde and 0.5-01% Saponin in PBS, as fixative and permeabilization medium.
6. The long-term storing of lentiviral VCM combined with the concentration and estimation MOI are regular procedure in many laboratories [18]. Often the procedure of virus concentration requires high-speed centrifugation, which is a difficult choice considering the obligations of BSL-2 working environment. It is same for the designated -80°C storing area. Therefore, we suggest the direct use of VCM without further processing. However, for the advanced usage, the desired MOI may not be obtained by this procedure. We demonstrate a simple method of commercially available polymer-based precipitation method using Lenti-X™ Concentrator, which can be replaced by in-house protocols using polyethylene glycol (PEG) precipitation or chondroitin sulfate combined with polybrene sedimentation. Similar consideration applies to retroviral preparations.
7. As described in the protocol, we use RetroNectin® coating optionally for enhancing the contact between lentivirus and the target cells. Similarly, we use poly-D-Lysine coating for settling the cells during retroviral transduction. However, these are optional methods. Simple centrifugation of cells together with viral supernatant supplemented with polybrene or protaminsulfate is sufficient. An alternative to these cationic polymers is peptide-based transduction facilitator such as Vectofusin-1 [19]. In our experience, these optional methods largely limit the cell death during transduction and only marginally change the transduction efficiency. To increase the transduction efficiency, we shall control the MOI, which best works between 2-5.
8. Lentiviral titer can be measured by several alternative methods, including p24 antigen, reverse transcriptase (RT) activity, and quantitative polymerase chain reaction modifications (qPCR). We suggest using the qPCR of Woodchuck Hepatitis virus posttranscriptional regulatory element (WPRE) region for estimating the titer, as described before [20].
9. We demonstrate the protocol with ecotropic retroviral delivery of sgRNA. Alternative to Phoenix Eco cells, ecotropic retrovirus could be prepared in HEK293T cells. Similarly, small sgRNAs can be introduced as oligo or as plasmid with reporter gene by transient transfection or electroporation [21]. For transient method, the analyses and sorting of the targeted cells need to be optimized within 2-4 days post transfection. This has advantage that we avoid continuous presence of the sgRNA post editing and therefore the gene function can be restored by cDNA without much consideration about the sgRNA target site.
10. Single cell sorting and preparation of clones is important for confirming the individual mutations introduced by Cas9 activity and checking clonal variations, if any. However, it is not necessary for assays involving the bulk input of the cells and for deletion of cell surface receptors that can be easily measured by FACS. Alternatively, the individual mutations can be determined in bulk by performing amplicon NGS using the same PCR product as used for sequencing of single cell clones. For intracellular targets, it is useful to perform single cell cloning to confirm the mutation as shown here (Fig. 3B). Sometimes, the targeted cells fail to develop single cell clones due to cell-cell contact and density dependent survival reasons. In that case, a two-step approach is a good alternative using 3-5 cells per well in the first round followed by single cell sorting.

## Acknowledgement

Authors thank Prof. C. Buske (CB) for critical reading of the MS and supporting the work. This work is supported by the Fritz Thyssen Stiftung (FTS) Grant 10.23.1.012MN to PCM, and Deutsche Forschungsgemeinschaft (DFG) CRC1279 project B01 to CB. The positions of MK and PCM are supported by FTS and DFG projects, respectively.

**Figure 3.**
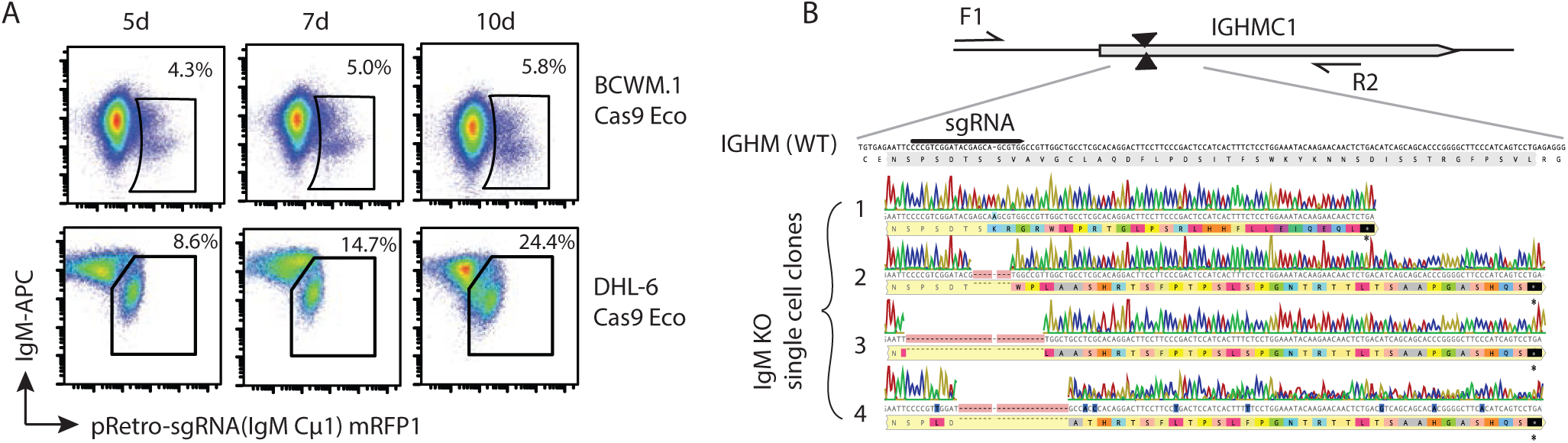
Generation of IgM deficient human lymphoma cell lines. A. Ecotropic retrovirus permissive B lymphoma cell lines BCWM.1 Cas9 Eco and DHL-6 Cas9 Eco were transduced with retroviral particle generated through pRetro gRNA (IgM) mRFP1 plasmid and FACS analyzed for mRFP1 and surface IgM expression by antibody staining after 5, 7 and 10 days post infection. Percent of cells within mRFP1 positive and IgM low gates were indicated within the plots. B. Representative sequence analyses of four IgM knockout (KO) BCWM.1 single cell clones analyzed by PCR amplification and sequencing of the 500 bp region encompassing the sgRNA target region as depicted in the figure.

