## Supplementary figures and images for "Facilitating Gene Editing in Human Lymphoma Cells Using Murine Ecotropic γ-Retroviruses"

### Supplemental Figure 1

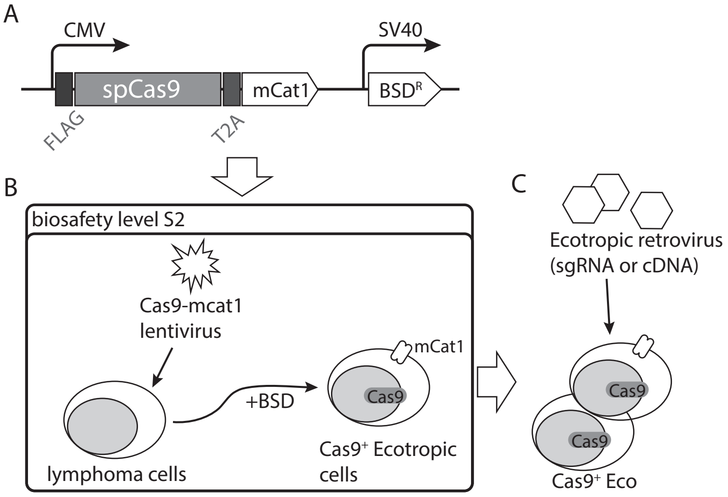

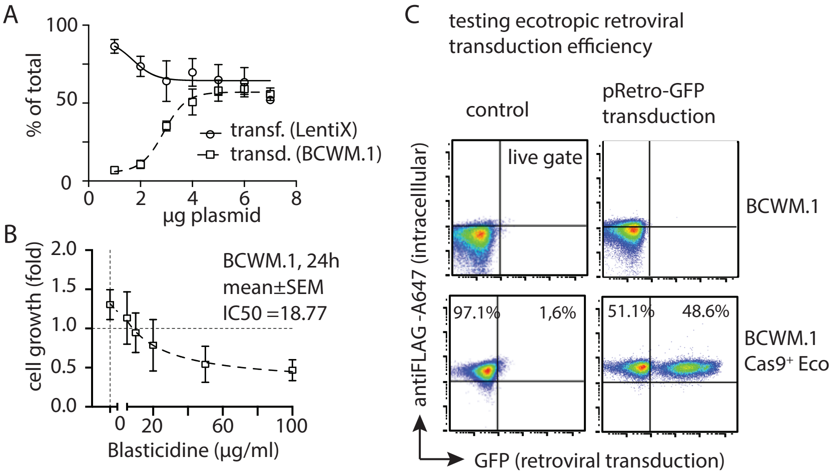

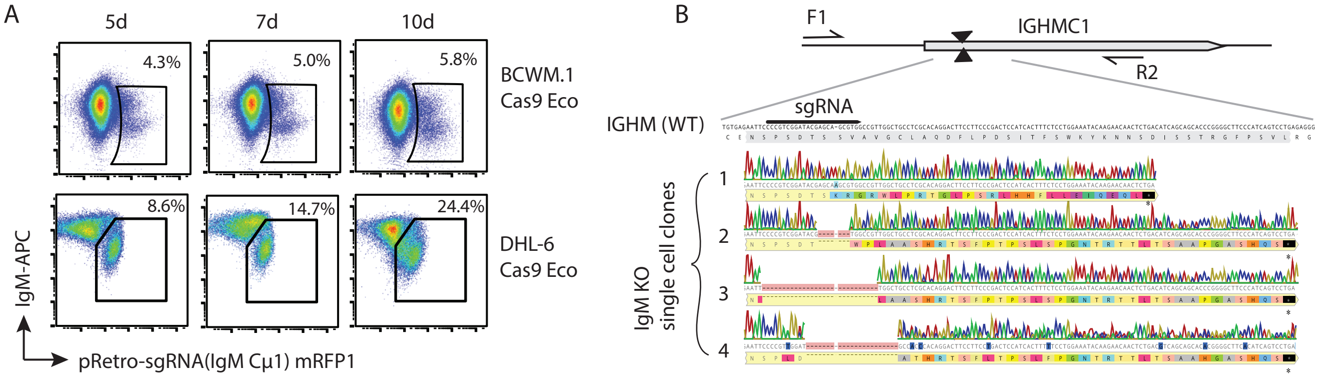
